# Inner speech and the neurobiology of psychosis

**DOI:** 10.1101/2025.08.22.671634

**Authors:** Jeremy I Skipper, Daniel R Lametti, David W Green

## Abstract

Aberrations of inner speech have been linked to psychotic symptoms such as thought insertion and auditory verbal hallucinations. These symptoms may reflect failures of prediction and source monitoring. Normally, efference copies of speech motor commands are sent to auditory cortices and suppressed, helping distinguish self-generated from external input. If suppression malfunctions, predicted auditory input may become perceptually salient. Further, if self-monitoring or error detection-related regions are also impaired (e.g., anterior cingulate cortex, ACC), inner speech may be misattributed as external. We tested this proposal using neuroimaging meta-analyses, examining how the brain systems in overt and inner speech production in neurotypical participants overlap with findings from psychosis-spectrum participants performing a range of tasks. They showed increased activity in motor-related regions associated with inner speech (e.g., ventral premotor cortices) and decreased grey matter in bilateral auditory cortices and ACC, in regions specific to overt speech. Coactivation-based network analyses revealed that these ventral premotor and auditory regions form distinct, inversely coupled audiomotor networks. Classification suggests the ventral premotor network supports ‘higher-level’ language processing, while the audiomotor network supports ‘lower-level’ speech and self-referential processing. Overall, results accord with the proposal that psychotic symptoms like auditory verbal hallucinations derive from phenotypic hyperactivation in inner speech-related regions that yield affectively salient efference copy signals that are insufficiently suppressed and monitored as self-produced. In line with a hierarchical predictive-processing account, disruption of a distributed recurrent system distorts self-awareness and conscious experience.

## Introduction

Inner speech, ‘the subjective experience of language in the absence of overt and audible articulation’^1^, goes by numerous names (e.g., ‘covert speech’ and ‘self-talk’) and is implicated in many psychological processes in both children and adults, comprising, for example, the regulation of emotions, behaviour, and cognition and reasoning about others (for review, see^1,2^). Arguably inner speech is also integral to, if not inseparable from, self-awareness and meta-self-awareness.^1–4^

Consistent with these important functional roles, inner speech occurs regularly, accounting for about 25% or more of all thought, though with considerable individual variability.^5–8^ Certainly, participants believe that inner speech occupies from about 50 to 75% of their thoughts,^9–14^ perhaps because inner speech, being central to so many fundamental psychological processes, including consciousness, is disproportionately salient in memory.^4^

Given its centrality in the lives of neurotypical individuals and its self-regulatory and evaluative functions, it is unsurprising that inner speech has also been implicated in a range of psychotic symptoms such as auditory verbal hallucinations (reported in up to 80% of patients), disorganized thinking, and thought insertion where a patient might state that ‘it feels like this person is using my brain to think his thoughts’.^13–29^ Good evidence for the relevance of inner speech to symptomology is increased lip muscle activity, consistent with subvocalization, observed in patients actually experiencing an auditory verbal hallucination aligned phenomenologically with their strong conviction as to its reality.^30–33^

What might be aberrant in the mechanism of inner speech in participants diagnosed with psychosis? As a starting point, consider the nature of typical overt speech. During production, corollary discharge or efference copy mechanisms (used here as if synonymous) allow the brain to predict the sensory consequences of self-produced speech, through feedback from motor to auditory cortices, suppressing responses to the predicted sounds, and allowing for rapid response and error correction.^21,22,29,34–39^ This account also requires that self-produced speech is distinguished from the speech of others. In speech production, this role is often attributed to anterior medial prefrontal brain regions like the anterior cingulate cortex (ACC) involved in the self/other distinction, source monitoring, and error detection.^40–46^ This overarching set of processes is not domain specific to speech production. Rather, it is general to most, if not all motor acts, e.g., the finger version of which famously accounts for why we cannot tickle ourselves.^47–51^ In inner speech, these processes have been suggested to work in much the same manner as in overt production.^4,52–54^ A motor command is issued, and an efference copy of that plan is sent to auditory cortex, dampening or suppressing neural responses to the predicted sounds.^4^ Such suppression normally prevents the utterance from being ‘heard’ but it also needs to be attributed to the self via processes associated with medial cortices like the ACC.^4^

Thus, the proposal for psychosis is that, if both attenuation and source-monitoring related functions of these distributed brain regions fail, the resulting sound of inner speech may be more vividly perceived and be misattributed to an external agent. Historically, efference-copy like theories and source-monitoring models of psychosis have been treated as largely separate domains, one focussing on ‘low-level’ sensory attenuation within speech circuitry, the other on ‘high-order’ tagging of source or ‘selfness’ involving the ACC.^55^ Contemporary predictive-processing frameworks instead situate these functions within a more expansive hierarchical model: efference copy conveys expected motor commands (subject to precision weighting as a function of overt or covert speech), while the ACC participates in evaluating prediction error and agency.^56–59^ In accord with this proposal, dysfunction can occur at least two levels: imprecise efference signals yielding unusually vivid inner speech, and under-weighted error signals fail to mark them as self-generated. Together, these produce the perceptual experience of an auditory verbal hallucination or thought insertion.

A predictive, efference copy account has been explored most extensively in the context of individuals experiencing auditory verbal hallucinations. Behaviourally, such individuals show evidence for deficits in sensory prediction and in distinguishing self-generated from external stimuli.^51,60–62^ Consistent with a breakdown in efference copy attenuation, they demonstrate reduced suppression of auditory responses during self-generated speech in EEG and MEG studies.^35,36,63,64^ Broadly, existing neuroimaging meta-analyses tend to support the involvement of brain regions and networks associated with inner speech^4,52,54,65–69^ and in line with a hierarchical model, specific studies report weakened connectivity between frontal speech-related regions and auditory cortex, along with reduced engagement of the ACC central to self-monitoring.^70–73^ Also noteworthy are reported changes in cerebellar connectivity in hallucinating patients given the cerebellum’s predictive role in speech.^74–76^ In contrast, a more recent meta-analysis suggests that the precise role of inner speech-related regions in the genesis of auditory verbal hallucinations remains unresolved in part because there were too few studies involving overt or covert inner speech production and self-referential processing to permit meta-analysis.^77^

Our aim in this paper was to establish such evidence meta-analytically, testing the hierarchical predictive coding model. Is there pathological perturbation in the brain systems involved in efference copy and source-monitoring mediating inner speech in the brains of psychoses spectrum participants? Critical to our approach, and, in contrast to prior meta-analyses, our hypothesis testing starts with a determination of the regions involved in overt and covert speech in neurotypical participants. First, we hypothesized that overt and covert speech engage overlapping but distinct regions in neurotypical individuals. Covert speech, lacking articulation, should rely less on the primary motor cortex and more on premotor regions. Overt speech involving actual movements and sound should rely more on the primary motor and auditory cortices. Second, given the proposed aberrations in efference copy mechanisms, we hypothesized that psychosis is associated with functional and structural alterations in the regions differentiating covert from overt speech. Third, we hypothesized that these regions form interacting networks reflecting the feedback connectivity expected in an efference copy mechanism. Finally, we hypothesized that regions involved in self-monitoring or source/error detection, such as the ACC, would also differentiate psychosis spectrum participants from neurotypical participants. Such a difference would help explain why aberrations in the efference copy mechanism are not recognized as internally generated.

To test these hypotheses, we conducted a series of neuroimaging activation and connectivity-based meta-analyses (the details of which are reported in the Methods). As noted above, our first step involved contrasting neuroimaging data for overt and covert speech production in neurotypical participants. Second, we identified functional and structural whole-brain differences with psychosis spectrum participants, analysed across a variety of tasks. Third, overlap analyses identified where psychosis-related differences intersected with the speech production regions and so provided a possible basis for inner speech dysfunction. At a novel fourth step, we performed a meta-analytic connectivity analysis to determine the network architecture of the speech production-related regions differentiating psychosis participants. Finally, we used meta-analytic functional profiling to characterize the psychological features most associated with the resulting networks.

## Results

### Neurotypical participants

Our goal was to better understand the relationship between inner speech production and symptoms such as auditory verbal hallucinations and thought insertion in participants diagnosed with psychosis spectrum disorder. Do the functional and structural differences in the brains of psychosis spectrum participants and neurotypical participants align with those from our hierarchical predictive coding model? To this end, we first contrasted overt and covert speech production in neurotypical individuals using ALE meta-analysis. We analysed 139 overt speech studies (4,590 activation locations) and 148 covert speech studies (3,935 locations) from the BrainMap database,^78–80^ with all results corrected for multiple comparisons using cluster-level family-wise error correction (P < 0.05). Overlap and contrast analyses were performed to identify shared and unique activation patterns between the two speech conditions.

As shown in Figure 1, covert and overt speech yield overlapping activity primarily in the left anterior insular cortex, inferior frontal gyrus (IFG) and sulcus (IFS), ventral precentral gyrus and sulcus, henceforth premotor cortex (PMv), and the posterior superior temporal sulcus at the location of the ‘temporal voice area’ or TVA^81^ (Fig. 1, top left, yellow). They also overlapped bilaterally in the medial superior frontal cortex at the location of the pre-supplementary motor area (pre-SMA; Fig. 1, bottom, yellow). Covert speech elicited more activity in the left IFG, PMv, inferior parietal cortex, primarily the intraparietal sulcus (IPS), and pre-SMA and the right pars opercularis of the IFG, among other regions. It lacked significant activity in ‘low-level’ audiomotor cortices (Fig. 1, blue). In contrast, overt speech was more associated with activity in these audiomotor regions. These included the bilateral central sulcus (M1/S1 or primary motor and somatosensory cortices), mid to anterior cingulate cortex (ACC), transverse temporal gyrus and sulcus (TTG or primary auditory cortex and TTS), and the nearby planum polare and planum temporale (immediately anterior and posterior to the TTG/TTS, respectively). Subcortically, overt speech production yielded more activity in the left putamen and bilateral thalamus and dorsal cerebellum (mostly lobule VI).

**Fig. 1.**
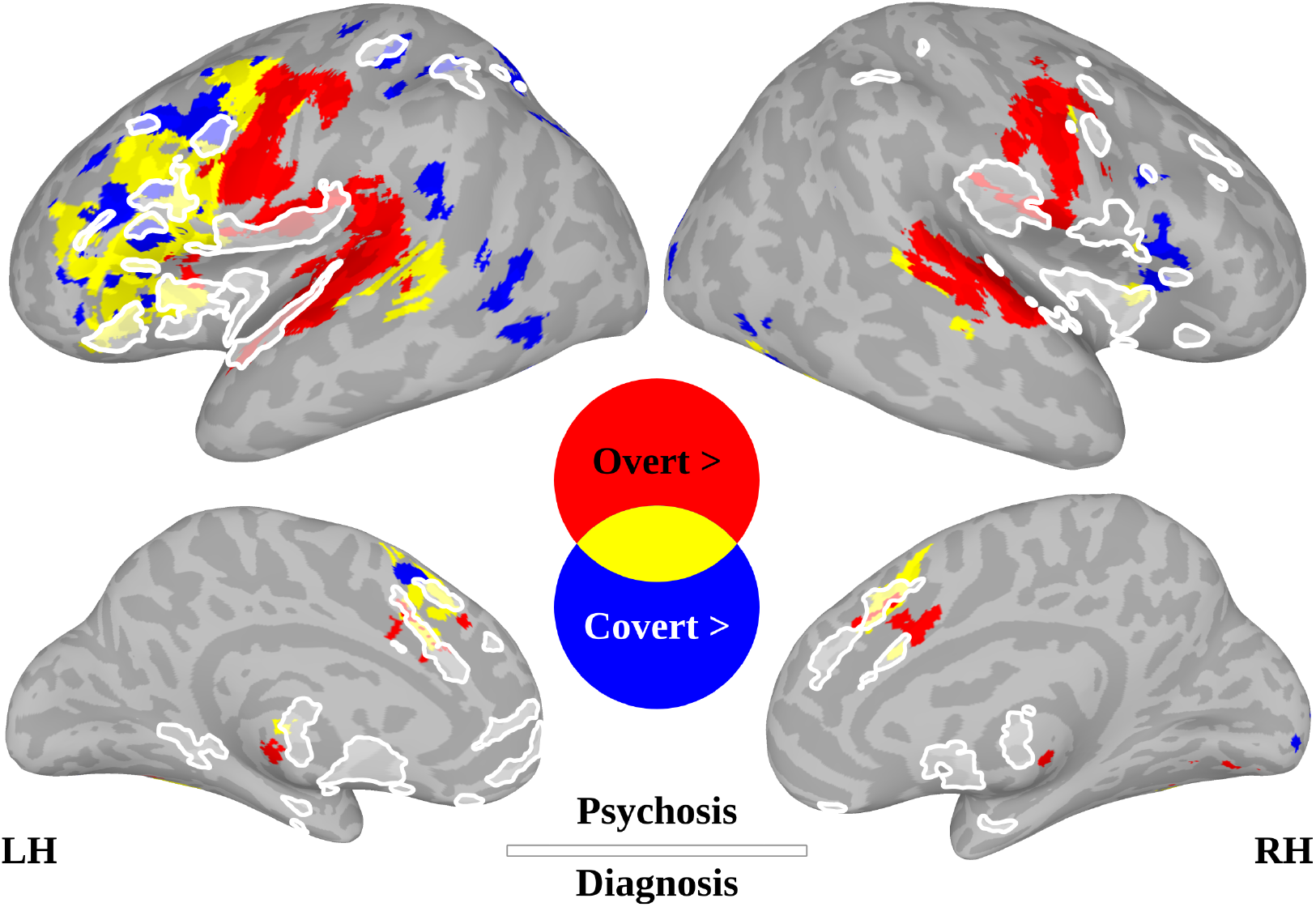
Meta-analysis of overt and covert speech production in neurotypical participants. Overt (red) and covert (blue) speech production were directly contrasted in neurotypical participants with region overlaps (yellow). To show the correspondence with speech, we provide a white outline of the functional and structural differences in psychosis spectrum diagnoses (which are shown in more detail in Fig. 2). All results are corrected for multiple comparisons using a family-wise error of 0.05, presented with a minimum cluster size of 100 voxels.

### Psychosis participants

We analysed 156 studies showing greater activation in psychosis spectrum participants than neurotypical participants (2,957 locations) and 104 studies showing the reverse pattern with activation greater in neurotypical participants than psychosis spectrum participants (1,378 locations). All studies were task-based and skewed towards emotional behaviours (with few overtly language-related studies), using predominantly face/visual picture-based paradigms (see Methods, Fig. 5). We also performed structural meta-analyses to determine grey and white matter differences, involving 48 studies showing greater grey/white matter in psychosis spectrum participants than neurotypical participants (389 locations) and 125 studies showing the converse pattern (1,760 locations; see Methods for the full breakdown and further information). Across all activation and structural studies, schizophrenia accounts for 77.74% of participants.

Figure 2 shows the results of ALE meta-analyses comparing functional and structural brain differences between individuals with a psychosis-spectrum diagnosis and neurotypical participants and overlaps with overt and covert speech production. Individuals with psychosis exhibited increased functional activity in the left anterior insula, bilateral IFG, PMv, and IPS. These regions primarily overlapped with those more engaged during covert speech production in neurotypical participants (Fig. 1, white outline and Fig. 2, red). Psychosis spectrum participants also exhibited decreased activity in the thalamus (Fig. 2, bottom left, blue). Structurally, psychosis spectrum participants showed decreased grey matter volume in the TTG (primary auditory cortex) bilaterally, which overlapped with regions more active during overt speech production in neurotypical participants (Fig. 1, red activity and white outline and Fig. 2, blue). Additionally, there were large decreases in grey/white matter volume throughout medial cortices that did not overlap with speech production, including even more anterior aspects of the ACC and orbitofrontal cortex. Subcortical patterns included decreased grey matter volume in the bilateral nucleus accumbens (NAc) and amygdala (Fig. 2, middle, blue), increased grey matter volume in the caudate and putamen (Fig. 2, bottom middle, red), and increased white matter in the thalamus and right cerebellum (Fig. 2, bottom right, red).

**Fig. 2.**
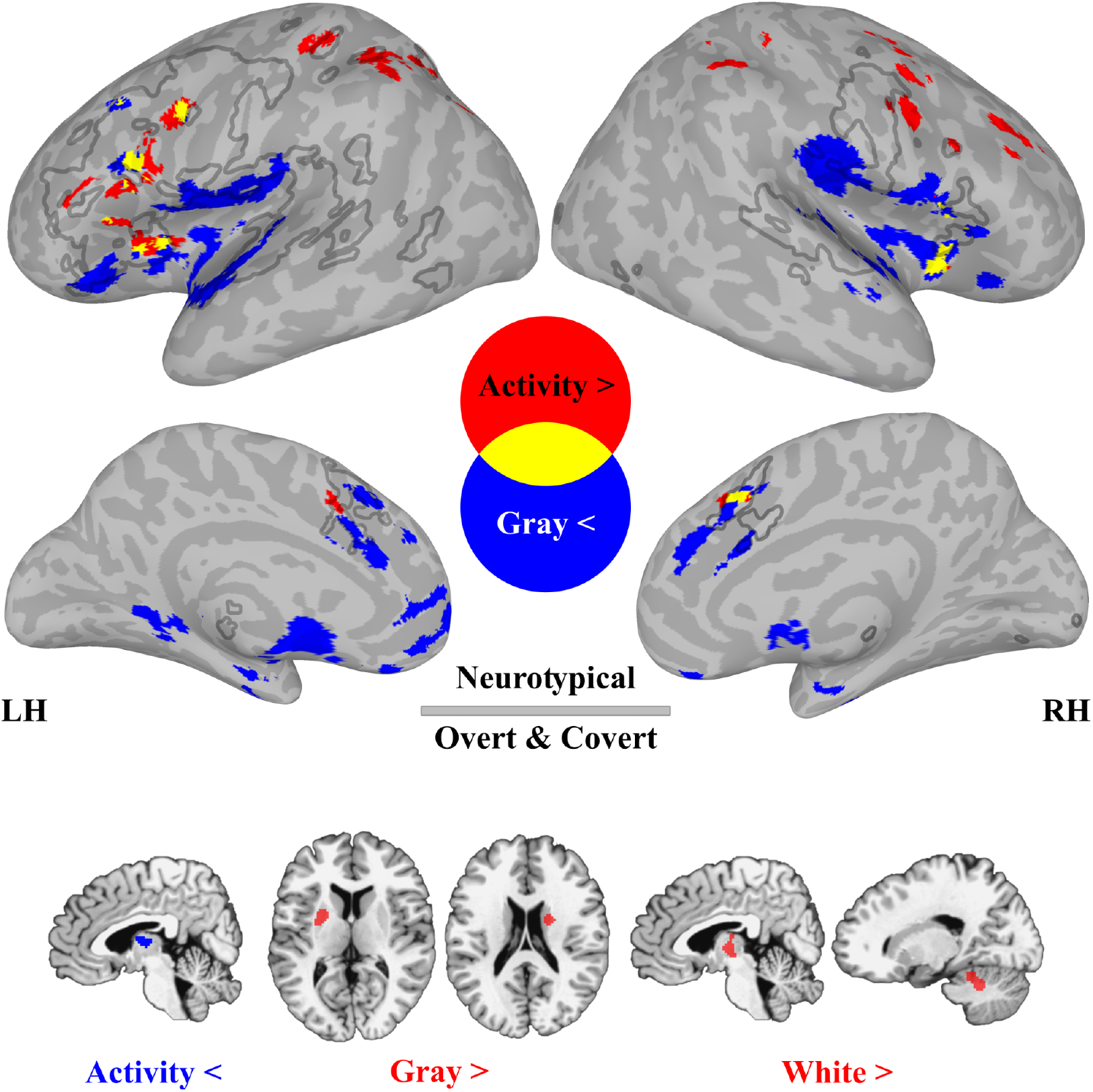
Meta-analyses of functional and structural differences in psychosis spectrum participants. There was significantly greater functional activity for psychosis spectrum diagnosis compared to the neurotypical controls (top surfaces, red) and a significant reduction in grey/white matter (top, blue). For comparison with Fig. 1, an outline of the contrasts and overlap of overt and covert speech production in neurotypical participants is shown (grey border). The remaining results on the bottom row (slices) were limited to subcortical structures, showing a decrease in thalamus activity, an increase in putamen and caudate grey matter and an increase in thalamus and cerebellar white matter for psychosis spectrum participants relative to neurotypical controls. All results are corrected for multiple comparisons using a family-wise error of 0.05, presented with a minimum cluster size of 100 voxels.

### Network analyses

To characterize the network organization of regions associated with speech production that differed in psychosis spectrum participants, we conducted a whole-brain coactivation-based meta-analysis using three clusters of interest identified from the overlap analysis of possible combinations of activity patterns (see Fig. 2). Specifically, the overlap clusters were generated using covert > overt and overt > covert speech production in neurotypical participants with psychosis spectrum participants > neurotypical participants, as well as the reverse contrast (neurotypical participants > psychosis spectrum participants) for both functional activation and grey/white matter data (using a minimum cluster size of 100 voxels). The first such cluster, located in the left pars opercularis (the most posterior aspect of the IFG) and PMv, showed increased functional activity in psychosis spectrum participants and overlapped with regions more active during covert speech production in neurotypical participants (Fig. 3, K1, black outline). The second and third clusters, located bilaterally in the TTG and TTS, showed reduced grey matter volume in psychosis spectrum participants and overlapped with regions more active during overt speech production (Fig. 3, K2 and K3, white outlines). These three clusters were then used in coactivation analyses of an independent set of 14,371 studies from Neurosynth.^82,83^ Coactivation maps show preferential coactivation with each individual cluster compared to other clusters, with each map corrected for multiple comparisons using false discovery rate (FDR) at q < 0.001.

**Fig. 3.**
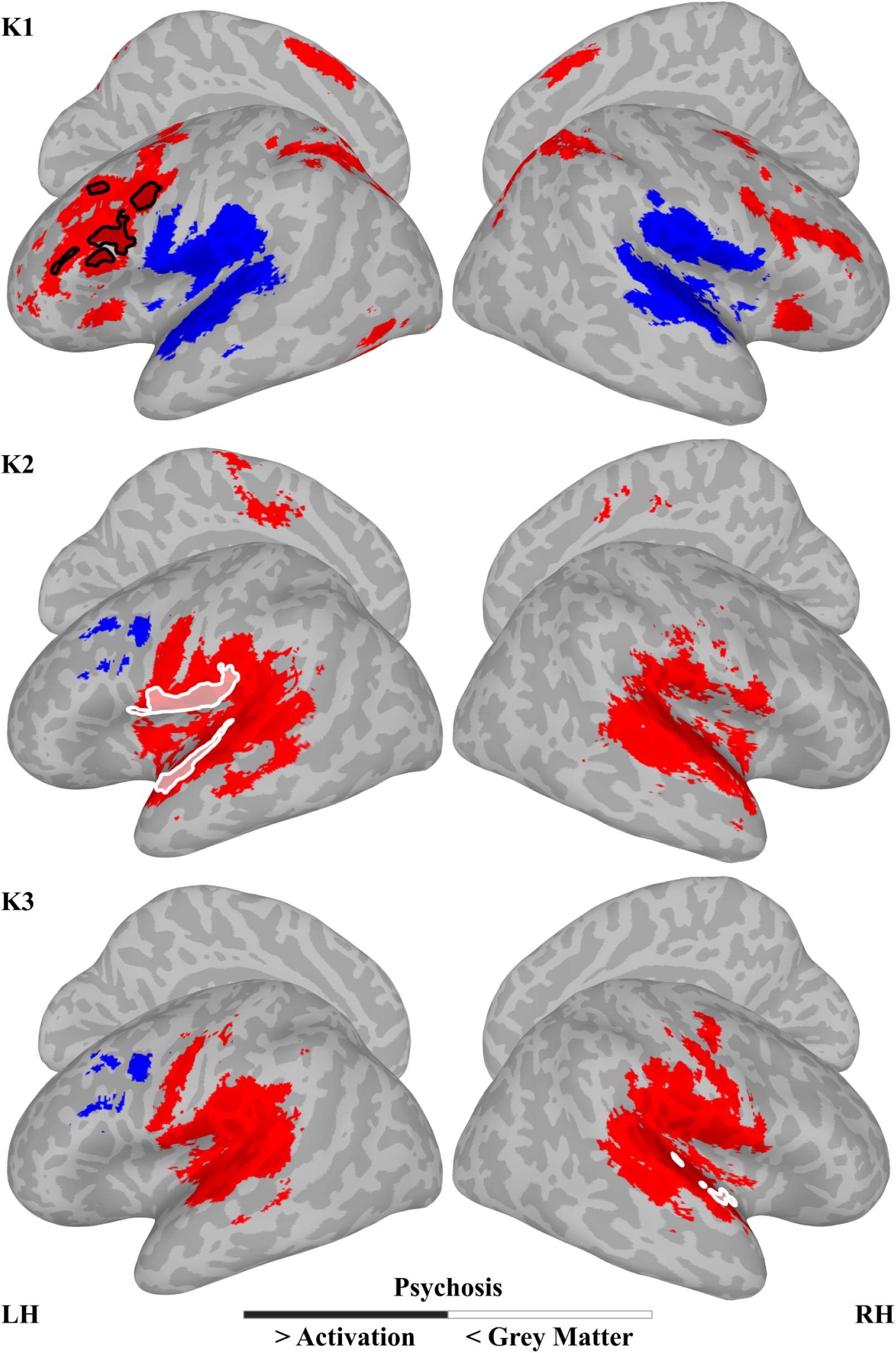
Meta-analytic connectivity from regions distinguishing psychosis spectrum from neurotypical participants in speech production-related regions. Three groups of clusters were used for coactivation analysis (K1-K3). The first included the left pars opercularis and ventral premotor cortex associated with greater functional activation in psychosis spectrum participants in regions more involved in covert speech production (black outline in K1). The other two included the left and right transverse temporal gyrus and sulcus associated with a reduction in grey matter volume for psychosis spectrum participants in regions more active during overt speech production (white outlines in K2 and K3). We identified the coactive (K1-K3, red) and co-deactivated (blue) regions for each of these clusters compared with the other two clusters across 14,371 studies. All results are FDR-corrected for multiple comparisons at q < 0.001.

Figure 3 shows that premotor regions (K1) and primary auditory regions (K2, K3) form distinct yet inversely coactive networks. Specifically, when one set of regions is more coactive across studies, regions resembling the others tend to be less coactive or deactivated. For example, K1 was associated with coactivation in a large portion of the ventral prefrontal cortex, including the inferior frontal gyrus bilaterally, as well as the left precentral gyrus and sulcus (i.e., PMv). Other coactivations included the anterior insula and more inferior parietal regions (like the IPS), bilaterally (Fig. 3, top, red). K1 also showed reduced coactivity in the TTG (primary auditory cortex) bilaterally, along with the planum polare and temporale, superior temporal gyrus (STG), operculum, and left ventral central sulcus (Fig. 3, top, blue). In contrast, K2 and K3 exhibited a near-opposite pattern, showing coactivation in bilateral TTG and surrounding auditory areas, as well as the ventral central sulcus (Fig. 3, bottom two rows, red), while producing less coactivation in the left ventral prefrontal regions (Fig. 3, bottom two rows, blue).

### Network profiles

To characterize the functional roles of the networks differing in psychosis spectrum participants, we analysed their broad activation profiles using large-scale meta-analytic functional decoding (Fig. 4). We employed the Neurosynth multivariate feature classification framework using a Gaussian Naive Bayes classifier, which achieved a mean accuracy of 67.33% in distinguishing the three clusters’ patterns of coactivity (K1-K3, Fig. 3). The network profile of these patterns was determined by their association with the top five features, i.e., meta-analytic maps with the largest z-transformed log-odds ratio (LOR). The left PMv cluster map (K1) was uniquely associated with ‘higher-level’ language terms, including the ‘words’ and ‘semantics’ meta-analyses (Fig. 4, red). In addition to these positive features, K1 was negatively associated with emotion, with three of the top five associated meta-analytic terms being ‘fear’, ‘pain’, and ‘reward’ (all LORs < −4.98 and P values < 0.000022). In contrast, the bilateral auditory cortex or TTG cluster maps (K2 and K3) were more strongly associated with ‘lower-level’ auditory and speech perception features, including the ‘auditory’ and ‘sound’ meta-analyses (Fig. 4, blue and white). Top-weighted negative features for this cluster encompassed self-other processing, including social cognitive (‘faces’ and ‘social’) and self-reference meta-analyses (all LORs < −4.03; P values < 0.00099).

**Fig. 4.**
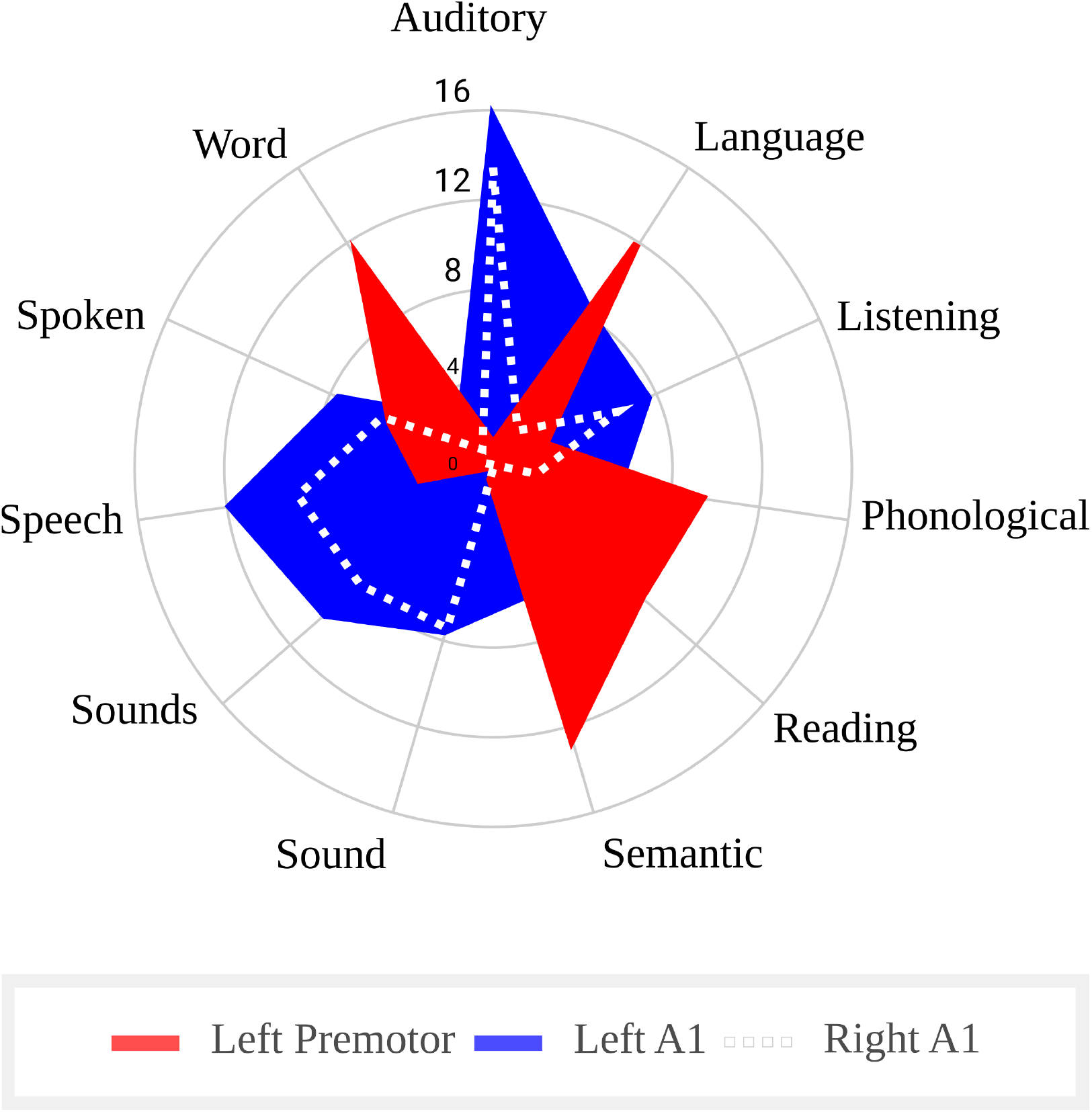
Functional profiles of network meta-analyses distinguishing psychosis spectrum and neurotypical participants in speech production regions. We used meta-analytically divided features to profile the three coactivation meta-analyses in Fig. 3. This resulted in 11 features, five features for K1 and six unique features for K2 and K3 (all LORs > 6.12; all FDR corrected P < 0.00000007). The radial plot displays the features most strongly positively associated with K1-K3 in Fig. 3. The coactivation map associated with left ventral premotor regions (K1 in Fig. 3) was more associated with ‘higher-level’ language features (red). In contrast, the coactivation maps associated with bilateral primary auditory cortices (A1, i.e., the transverse temporal gyrus/sulcus; K2 and K3 in Fig. 3) were more associated with ‘lower-level’ auditory/speech features (blue and white dots).

**Fig. 5.**
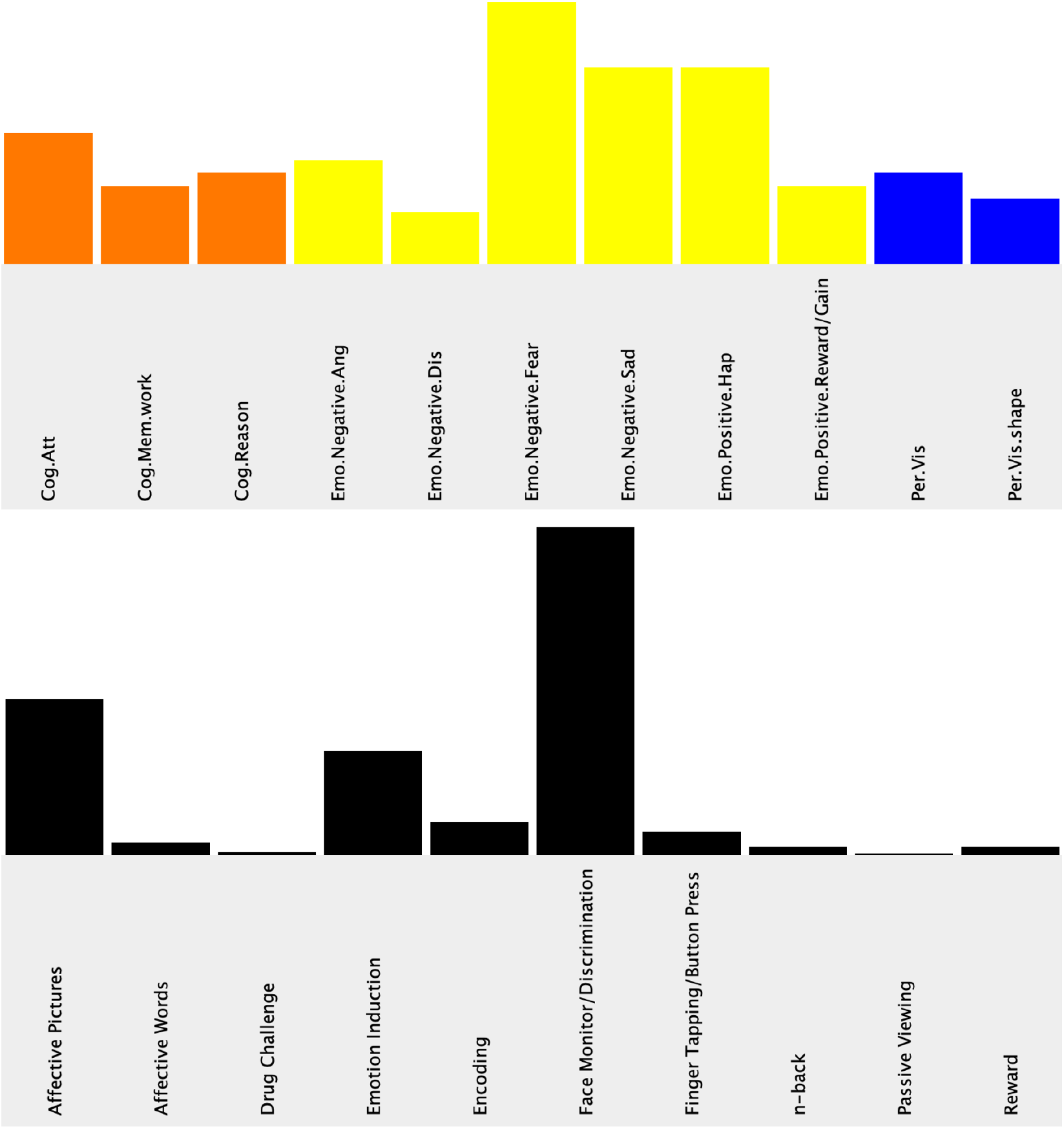
Behaviours and paradigms in included studies. The top row (coloured bars) is the behavioural histogram, and the bottom row (black bars) is the paradigm class histogram reported by BrainMap for the studies used in the manuscript.

## Discussion

We tested a unified account of inner speech dysfunction in psychosis spectrum participants. It situates dysfunction at two levels within a predictive coding hierarchy: lower-level motor-sensory loops generate and attenuate predictions, while higher-order regions such as the ACC regulate the precision of prediction errors to support agency attribution. Our coordinate-based neuroimaging meta-analyses largely confirm and extend this account.

We identified specific patterns of brain differences in psychosis spectrum participants that overlap with regions distinguishing overt from covert speech production in neurotypical participants (Fig. 1). Hyperactivity was observed predominantly in regions where covert speech production exceeds or overlaps with overt speech, including ventral premotor cortex, inferior frontal gyrus, anterior insula, thalamus, putamen, and cerebellum, largely avoiding regions more strongly engaged during overt speech (Fig. 2, white outline; Fig. 3, red). In contrast, grey matter volume reductions occurred in bilateral primary auditory cortices, medial prefrontal cortex, including the ACC, primarily overlapping regions where overt speech production exceeds covert speech (Fig. 2, white outline; Fig. 3, blue). This functional-structural dissociation extended to regions not traditionally associated with speech production, including the NAc, amygdala, orbitofrontal cortex, and additional medial prefrontal cortices, including more extensive aspects of the ACC. Network analyses demonstrated that these anatomically distinct sets of regions form inversely coupled networks (Fig. 3). Overactive premotor regions were more coactive with prefrontal and parietal regions and associated with ‘higher-level’ language functions, while structurally reduced auditory regions were more coactive with audiomotor regions and associated with ‘lower-level’ speech processing (Fig. 4), both negatively associated with emotions and self-referential processing.

Taken together, these results suggest the following view of the aberrant nature of inner speech in psychosis spectrum disorders. Psychosis spectrum disorder is associated with hyperactivation of speech-planning-related brain regions involved in inner speech. These include regions critically implicated in efference copy mechanisms, such as the ventral premotor cortex, involved in planning and controlling laryngeal, lingual, and labial articulators, along with the cerebellum, a key structure in predictive processing in both speech perception and production.^76^ Further, these regions are associated with higher-level language functions, consistent with their role in using contextually derived language representations as hypotheses to predict speech sounds arriving in auditory cortices through efference copy.^84–86^ The observed reductions in grey matter volume in primary auditory cortices suggest a compromised capacity for sensory processing and inhibition of predicted auditory feedback. This accords with findings that progressive grey matter loss in temporal (auditory) and frontal regions is associated with the emergence and severity of auditory hallucinations and psychotic symptoms. Structural changes may then underlie deficits in predictive and feedback processing.^87–93^ The inverse coupling of networks associated with the hyperactive premotor and aberrant auditory regions is what might be expected from a predictive system involving feedback in which one set of regions precedes processing in another. Finally, the dramatic reductions in grey matter volume in regions typically associated with emotional regulation and self-monitoring (e.g., the ACC) suggest that aberrant inner speech processing may be less readily recognized as being self-produced.

Our results support an integrated structural-functional model of dysfunctional inner speech leading to symptoms like auditory verbal hallucinations or thought insertion. We have already described the first three components of this model that accord with dysfunction in two levels of a predictive coding account. First, hyperactivation in motor speech regions generates excessive and/or stronger efference copy signals during inner speech. Second, structural compromise in primary auditory cortices impairs the normal suppression of these predictive signals. Third, structural reductions in auditory cortices, combined with the ACC, disrupt source monitoring processes that normally attribute inner speech to the self. The structural reductions may represent developmental abnormalities or progressive changes that necessitate compensatory hyperactivation in preserved motor regions. However, this compensation is ultimately maladaptive, creating an imbalance between prediction generation and suppression that manifests as hallucinated voices or thought insertion.

Our other findings suggest extensions of this model. Various changes in the anterior insula, NAc, amygdala, orbitofrontal cortex, and medial prefrontal regions, including the ACC, allow us to envisage integration with research on salience and dopamine dysregulation.^94^ That is, these regions may reflect the pathological salience attributed to internally generated speech that has escaped normal predictive suppression, given its vividness and affective significance. A plausible mediator of such amplification is elevated mesolimbic dopamine, enabling the enumerated regions we found to imbue predictions with motivational and affective significance.^94,95^ Dopaminergic amplification may therefore be a contributor to a larger set of mechanisms involving efference copy and source monitoring rather than a separate mechanism per se. Finally, thalamic and basal ganglia differences suggest broader alterations in the circuits that gate and filter sensory information and may serve an integrative role, linking the distributed systems involved in inner speech, efference copy, source monitoring, dopamine, and salience, thereby cumulatively reflecting failures of predictive suppression mechanisms.

We note two constraints on interpretation. First, although the heterogeneity of psychosis-spectrum disorders and paradigms included in our analyses allowed us to identify a major phenotypic difference in psychosis spectrum participants relative to neurotypical participants (by increasing sample size), it may have obscured disorder- and task-specific patterns of predictive processing dysfunction. However, greater than 75% of participants had schizophrenia, and most tasks were not language oriented, likely minimising this concern. Second, the cross-sectional nature of the included studies prevents causal inferences about the source of the differences (functional and structural) with neurotypical participants. Longitudinal studies tracking individuals from prodromal stages through the first episode would be needed to establish causality. Lastly, we acknowledge that layer-specific imaging will likely be necessary to assess whether or not predictive and error-correction work as proposed in our predictive coding model.^57^

These limitations aside, our findings and proposed model have important implications for both the treatment of psychosis and theories of consciousness. They identify specific neural targets (e.g., hyperactive premotor regions, reduced auditory and medial frontal cortices) for potential therapeutic interventions. Indeed, transcranial magnetic stimulation targeting these regions has shown promise in reducing auditory verbal hallucinations,^96–98^ while real-time fMRI neurofeedback aimed at recalibrating predictive loops may offer even more precise interventions.^99–102^ More broadly, our results illuminate the foundations of conscious experience. The predictive processing model of psychosis we propose suggests that consciousness depends on the integrity of recurrent prediction–error loops linking motor and sensory systems, in line with recurrence-based theories of consciousness and the central role of prediction in perception and awareness.^103^ They also provide neural grounding for accounts that position inner speech as fundamental to self-awareness and meta-self-awareness, converging with global neuronal workspace and higher-order thought theories.^4,104,105^ From this perspective, psychosis represents not merely a clinical syndrome but a natural perturbation of the recurrent predictive mechanisms that normally sustain self-awareness and conscious experience.

## Methods

### Database search

We conducted neuroimaging meta-analyses using the BrainMap database (http://brainmap.org/, accessed December 2020). We programmatically queried the database for studies reporting stereotaxic coordinates (x, y, z) of functional brain activity based on specific metadata criteria.^78–80^ All coordinates originally reported in Talairach space were converted to Montreal Neurological Institute (MNI) space using established transformation algorithms to ensure spatial consistency across studies.^106,107^

In order to establish the brain regions involved in speech production, we performed searches for overt and covert speech in neurotypical participants. To maximise statistical power and study inclusion, no restrictions were applied for handedness or age, since such constraints reduced the number of eligible studies. The search criteria included studies with normal mapping context, activation-only results, and paradigm classes that included both overt and covert variants. Specifically, in pseudocode (where “&” = “and”; “|” = “or”), the search criteria used were:

> [Experiments, Context, is, Normal Mapping] & [Experiments, Activation, is, Activations Only] & [Experiments, Paradigm Class, is, Naming (Overt | Covert) | Reading (Overt | Covert) | Recitation/Repetition (Overt | Covert) | Word Generation (Overt | Covert) | Word Stem Completion (Overt | Covert)]

The overt speech search yielded 139 studies (1,942/2,317 participants, 469/874 experiments, 510/595 conditions, 4,590/8,125 locations), while the covert speech search returned 148 studies (2,039/2,411 participants, 441/799 experiments, 498/630 conditions, 3,935/6,903 locations).

### Psychosis studies

For psychosis-related analyses, initial searches using the same speech-specific paradigm classes with psychosis diagnoses yielded insufficient studies for robust meta-analysis. Applying age and handedness restrictions resulted in no studies, while removing these restrictions yielded only nine overt and six covert studies (approximately 120 locations each). We therefore broadened our inclusion to all functional activation studies involving participants with psychosis-spectrum diagnoses, based on the hypothesis that alterations related to inner speech, if fundamental to the phenotype, would manifest across diverse experimental paradigms. The expanded search criteria included activation-only studies covering seven diagnoses. The search conducted was:

> [Experiments, Activation, is, Activations Only] & [Subjects, Diagnosis, is, Family History of Schizophrenia | Nonschizophrenia Psychosis | Psychosis | Psychotic Major Depression | Schizoaffective Disorder | Schizophrenia | Schizotypal Personality Disorder]

This broader search returned 194 studies (7,301 participants, 1,161 experiments, 592 conditions, 8,165 locations).

Manual curation was performed to generate two datasets, with activation foci specific for psychosis-spectrum participants (patients) and neurotypical participants (controls). We extracted only results involving participants with a psychosis diagnosis, healthy controls, or explicit patient-control contrasts (patients > controls or controls > patients). Inclusion criteria were patients > controls contrasts, patients-only activations, clear directional patient versus control comparisons with appropriate statistical values, and patient-control conjunctions. Exclusion criteria included main effects and interactions that were not clearly patient-specific and bidirectional contrasts without clear directionality to indicate whether activity was more for patients or controls. This process yielded 156 studies for patients > controls (4,385 participants, 461 experiments, 447 conditions, 2,957 locations) and 104 studies for controls > patients (3,976 participants, 238 experiments, 279 conditions, 1,378 locations). Fig. 5 provides an overview of the behaviours and paradigms involved in the activation studies in the resulting datasets.

### Structural meta-analyses

To assess corresponding structural brain alterations, we performed additional searches for grey and white matter differences between psychosis-spectrum participants (patients) and neurotypical participants (controls). Search criteria included Grey Matter or White Matter contrasts, Controls > Patients or Patients > Controls contrasts, and psychosis-spectrum diagnoses (Family History of Schizophrenia, Nonschizophrenia Psychosis, Psychosis, Schizoaffective Disorder, Schizophrenia, Schizotypal Personality Disorder, Subclinical Psychosis). Note that the functional search included

Psychotic Major Depression but not Subclinical Psychosis, while the structural search included Subclinical Psychosis but not Psychotic Major Depression. The specific search criteria in pseudocode for the two meta-analyses were:

> [Experiments, Contrast, is, Gray Matter | White Matter] & [Experiments, Observed Changes, is, Controls > Patients | Patients > Controls] & [Subjects, Diagnosis, is, Family History of Schizophrenia | Nonschizophrenia Psychosis | Psychosis | Schizoaffective Disorder | Schizophrenia | Schizotypal Personality Disorder | Subclinical Psychosis]

This yielded four datasets: patients > controls grey matter (39 studies, 3,115/3,950 participants, 49/177 experiments, 335/1,207 locations), controls > patients grey matter (94 studies, 7,893/8,776 participants, 138/321 experiments, 1,554/2,556 locations), patients > controls white matter (9 studies, 767/860 participants, 13/55 experiments, 54/322 locations), and controls > patients white matter (31 studies, 2,470/2,688 participants, 42/125 experiments, 206/677 locations).

### ALE meta-analyses

Coordinate-based meta-analyses were performed using the Activation Likelihood Estimation (ALE) algorithm.^108–111^ Each MNI coordinate was modelled as a three-dimensional Gaussian probability distribution, and ALE values were computed to quantify the convergence of reported activations across studies. We used the union of these distributions to compute an ALE map, which quantifies the statistical convergence of activation foci across experiments. We assessed statistical significance via permutation testing (1,000 permutations) to evaluate above-chance spatial clustering. All resulting ALE maps were corrected for multiple comparisons using a cluster-level family-wise error (FWE) correction of P < 0.05, with an individual voxel-wise threshold of P < 0.001. Contrasts and conjunctions between ALE maps were computed using the same permutation-based procedure and correction parameters, with a minimum cluster size of 200 mm3 for significance and 100 voxels for display purposes.

### Overlap analyses

To identify regions at the intersection of speech production networks and psychosis-related alterations, we performed overlap analyses using the 3dcalc utility in AFNI.^112^ Complex Boolean expressions were employed to delineate unique and shared regions of activation or structural alteration, isolating voxels showing overt > covert speech activation overlapping with psychosis-related changes, covert > overt speech activation overlapping with psychosis-related changes, and shared activation patterns between conditions. Network meta-analyses further examined the intersection of speech-related and psychosis-related clusters. Results were thresholded at P < 0.05 with minimum cluster sizes of 100 voxels for visualization, as shown in Figs. 1–3.

### Coactivation meta-analyses

The functional connectivity of the three key clusters identified in the overlap analysis (Fig. 3) was investigated using coactivation meta-analyses within the Neurosynth framework, which leverages a dataset of 14,371 studies.^82,83^ Identified clusters from the overlap analyses were resampled to Neurosynth MNI template space. Each cluster was used as a seed region to generate a whole-brain coactivation map. Coactivation contrasts were computed to identify brain regions that preferentially coactivated with each cluster relative to others, with results corrected for multiple comparisons using FDR at q < 0.001.

### Machine learning classification

To functionally characterize the identified networks, we employed the Neurosynth multivariate feature classification framework.^83^ A Gaussian Naive Bayes classifier was trained to predict which of the three seed clusters was associated with a given study’s activation pattern, with classification accuracy evaluated using cross-validation. The classifier achieved a mean accuracy of 67.3% (individual region accuracies: 64.3%, 69.0%, 68.7%). Feature importance was quantified using log-odds ratios, and the top five discriminative psychological terms (excluding anatomical terms) for each network were visualised on polar plots with reordering based on importance scores, as presented in Figure 4.

### Statistical software

All ALE analyses were conducted using GingerALE.^79,80^ Image calculations and manipulations were performed with AFNI (Analysis of Functional NeuroImages).^112^ Coactivation and classification analyses utilized Python-based Neurosynth and scikit-learn machine-learning libraries.^82,83^ Surface-based visualisations were created using SUMA (Surface Mapping) with custom colour palettes and cluster outlines for publication-quality figures.^113^ All final statistical maps and overlays were presented with the specified correction for multiple comparisons and minimum cluster-size thresholds.

## Acknowledgements

We thank Rachel Morse and the LAB Lab and UNITy Project more generally for discussion. JIS is supported by the Wellcome Leap and BB.

## Author contributions

JIS conceived of the study and did all the analyses, made all figures, and drafted the initial manuscript. DG and DL helped conceptualise and redraft.

## Open science statement

This study was not preregistered. A version of this paper will be made publicly available as a preprint on bioRxiv. The code used to generate results will be made available on GitHub (https://github.com/lab-lab/).

